# Gel-Amin for improving signal propagation and extracellular recordings of cardiomyocytes in a 3D Microphysiological System

**DOI:** 10.1101/2025.09.04.674300

**Authors:** Dominic Pizzarella, Katelyn Neuman, Nolan Burson, Abigail N. Koppes, Ryan A. Koppes

## Abstract

Microphysiological systems (MPSs) hold great potential for fundamental discovery and accelerating the drug discovery pipeline through simplifying complex tissues to their first principles and enabling real-time, high-resolution monitoring. Hydrophilic biomaterials, such as hydrogels, are important for MPS innovations due to their ability to emulate the native extracellular matrix and tunable mechanical properties. Furthermore, hydrogels can be tailored to improve tissue maturity as well as the efficacy of instrumentation. However, many biopolymers are non-conductive, presenting complications for modeling excitable tissue environments like the heart. In this work, we show that an 8% (w/v) Gelatin Methacryloyl (GelMA) + 3.5% (v/v) Choline Acrylate hydrogel, nicknamed Gel-Amin, can amplify extracellular voltage recordings from a culture of cardiomyocytes (CMs) from commercial microelectrode arrays. Our laser cut and assemble method for manufacturing 3D MPSs allowed direct comparisons of CM signal propagation in Gel-Amin compared to control GelMA cultures in a single system. This innovative material supported *in vitro* CM cultures with improved synchronicity and greater signal-to-noise ratios (SNRs), suggesting potential improvements over conventional biomaterial limitations. Here, we developed a cost-effective *in vitro* cardiac tissue model that allows real-time electrical activity monitoring.

## Introduction

Cardiotoxicity, damage to the heart caused by a therapeutic agent, is one of the most frequent adverse drug reactions in late-phase clinical trials.^1-3^ Drug development is quite inefficient, costing $1-2 billion and spanning an average of 10-15 years before Food and Drug Administration (FDA) approval.^4, 5^ Three-dimensional (3D) *in vitro* microphysiological systems (MPSs)^6-8^ have shown promise in recapitulating key features of human physiology, including the cardiovascular system^9^ to offer insights into disease mechanisms and tissue organization. A leading application of these *in vitro* cardiac platforms is the drug discovery pipeline to evaluate preclinical compounds on human biology. However, finding the balance between simplicity, measurable biomarkers, and appropriate biorelevancy remains a challenge. Planar microelectrode arrays (MEAs) are relatively user-friendly instruments for capturing high-resolution extracellular recordings of cardiomyocyte (CM) action potentials (APs), yield direct insight into real-time cell health in response to external stimuli.^10, 11^ Utilizing these platforms for 3D tissue culture remains a challenge as signal resolution relies on cell proximity to the recording electrode.^12^ This requires custom array fabrication^13^, conformable MEAs^14^, or 3D penetrating electrodes^15^.

Within MPSs, biomaterials are often composed of natural polymers, such as collagen, gelatin^16^, and alginate, or synthetic polymers, like poly(lactic acid) (PLA), and have commonly been employed to encapsulate primary cell cultures in 3D hydrogel constructs.^17^ Many of polymeric materials are advantageous for their tunable mechanical properties and cell adhesion properties. To develop mature cardiac tissue *in vitro*, hydrogels that facilitate signal propagation are needed to encourage coordinated contraction in 3D engineered tissues.^18^ The ideal cardiac tissue engineering scaffold would match the native heart’s mechanical properties, mimic the microenvironment of the cardiac system, promote cardiac cell viability, cell attachment (biocompatibility), and have the appropriate conductivity (conductivity range of 300 × 10^−5^ S cm^−1^ to 600 × 10^−5^ S cm^−1^ for native myocardium) to allow propagation of electrical stimulation needed for cardiac cell communication.^19, 20^

Increasing bulk electronic conductivity of biomaterials has traditionally been accomplished through metallic, carbon, or PEDOT:PSS doping^21^. While conductivity is increased by electron transport in these dopants, it is often at the cost of optical transparency, rendering live and endpoint microscopic-based analyses nearly impossible.^22^ Alternatively, bio-ionic liquids (Bio-ILs) are nontoxic, biodegradable organic salts that are liquid at room temperature and have ionic conductivity and electrochemical stability.^23^ Choline acrylate (ChoA) has been a recent Bio-IL of interest in tissue engineering due to choline’s biocompatibility.^24^ Choline is an essential micronutrient involved in many physiological functions and can be readily broken down for reuse.^25^ We have recently investigated ChoA conjugated to GelMA (Gel-Amin) for peripheral nerve repair applications.^26, 27^ Gel-Amin exhibits higher conductivity of a collagen-based hydrogel without adverse effects on neuron cell viability or optical transparency.^26, 27^

We have previously developed a cut & assemble method for fabricating 3D MPSs.^9, 28-31^ This method of fabricating organs-on-a-chip lends itself to incorporating instrumentation, including MEAs for real-time recording of CM contractions. Furthermore, our GelPin technology allows contiguous interfacing of two unique hydrogel compositions to investigate signal propagation between collagen-based biomaterials of different conductivities within one platform. Here, we utilize our 3D MPS to evaluate the impact on CM signal propagation through interfacing GelMA and Gel-Amin (**Fig. 1**). We hypothesized that the inclusion of Gel-Amin would improve the ionic conductivity, enhancing AP propagation to recapitulate coordinated cardiac contractions and SNR of recording electrodes. To study this, the group developed a high-throughput, accessible, and cost-effective MPS capable of recording CM signaling via a MEA.

**Figure 1.**
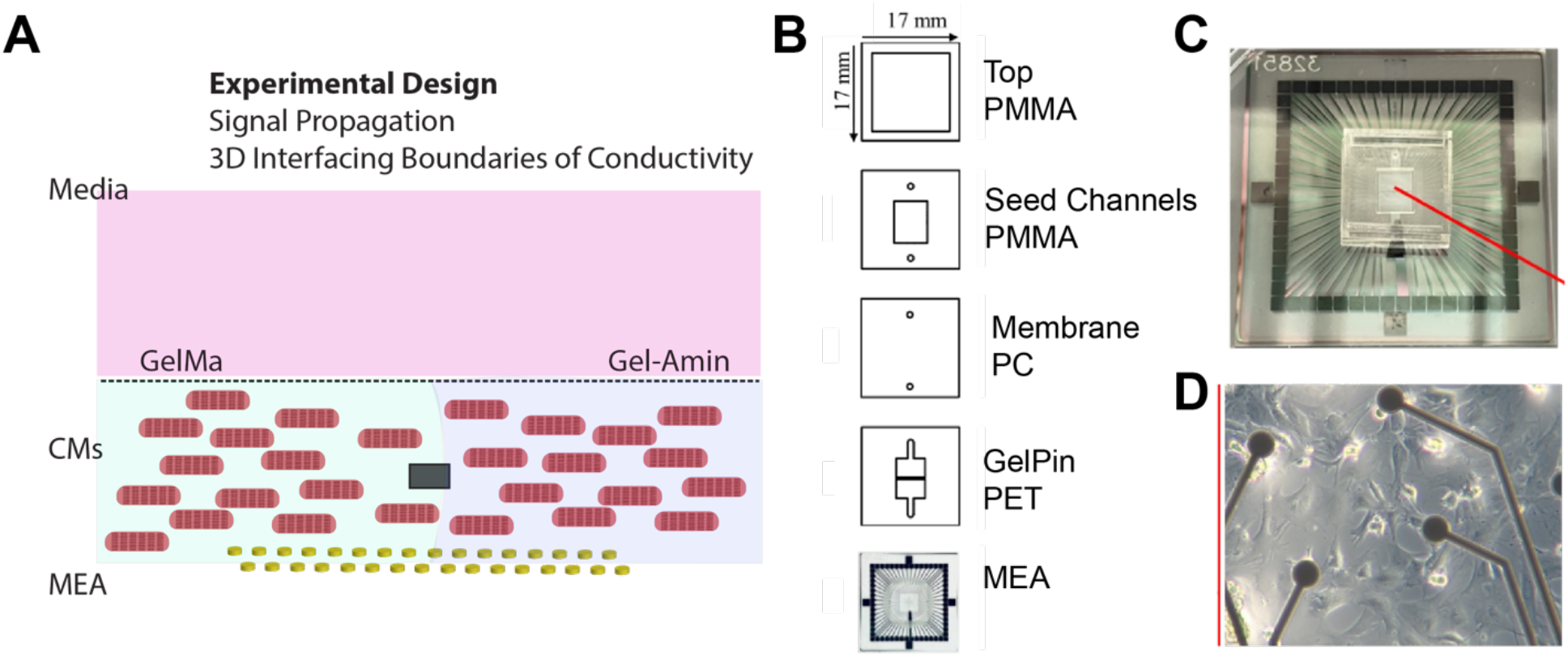
3D MPS platform to interface CM-laden collagen-based biomaterials of different electrical conductivities. A) Encapsulated cardiomyocytes are contiguous yet constrained within their contrasting hydrogels via a floating GelPin layer. B) MEA Chip Design. I) The top channel was optimized to hold and exchange media. This region was comprised of 3/16” PMMA with one layer of 3M tape. II) The seed channel was made from 1/16” PMMA and allowed for a controlled seeding region. One layer of 3M tape was used to stick it to the membrane. III) A 1.0 μm pore membrane. IV) The GelPin layer was made from 0.005” PET with two sides of 3M tape. One side was connected to the membrane, and the other was attached to the glass MEA. The GelPin was designed to separate the two hydrogel groups from one another on one MEA. IV) MEA was at the bottom of the design. C) Chips were positioned to equally separate the electrodes. D) The assembled MEA Chip is shown with a representative image of cells encapsulated in the Gel-Amin hydrogel. (electrode diameter = 30 μm)

## Methods

### Hydrogel Fabrication and Characterization

Both GelMA and Gel-Amin batches were synthesized and characterized as previously reported by Neuman et al.^27^ In brief, hydrogel precursor solutions were prepared to contain 0.5% (w/v) lithium phenyl-2,4,6-trimethylbenzoylphospinate (LAP; Allevi) photoinitiator. The precursor solution was then photocrosslinked with blue light (405 nm, 10 W). Exposure time was calculated as a function of the height of the hydrogel (0.25 s of exposure time per µm).

Hydrogel samples were mechanically evaluated with a TA Instruments Electroforce 3200 universal mechanical testing platform. During compression testing, samples were submerged in DPBS to preserve a hydrated environment. Dynamic Mechanical Analysis (DMA) was used to perform a compressive, sinusoidal frequency sweep between 0.5 and 5 Hz with 10% strain. DMA analysis was performed with WinTest® 7 software. A Fourier analysis on the data collected was performed to calculate the difference in phase (*δ*) between the dynamic peak-to-peak force function and the dynamic peak-to-peak displacement amplitude. δ was then used to calculate the storage modulus (*E*′), loss modulus (*E*′′), and complex modulus (*E*\*).

Electrochemical Impedance Spectroscopy (EIS) of GelMA and Gel-Amin hydrogels was performed using a VSP Potentiostat (BioLogic). Cylindrical hydrogels (Ø = 8 mm, height = 7 mm) were cut to a uniform size with a 6 mm diameter biopsy punch (Fisher Scientific) and inserted into a custom-designed cell made of perfluoro alkoxy alkane (PFA; Grainger). Magnesium stick electrodes (Lincoln® Electric) were inserted into the cell to be in contact with the sample. Dulbecco’s Phosphate-Buffered Saline (DPBS) was pipetted into the cell and surrounded the sample. EIS was performed from 1 MHz to 100 mHz with a sinusoidal amplitude of ±10 mV. Data analysis was performed with EC-Lab® Software.

### Primary cell isolation

Primary CMs were isolated from two-day-old (p2) Sprague–Dawley neonatal rats^9,32^ as approved by Northeastern University’s Institutional Animal Care and Use Committee. Briefly, hearts were extracted from rats and stored in a 0.05% trypsin (v/v in Hanks’ Balanced Salt Solution; HBSS) solution for 16 hours at 4°C. Following this, cardiac tissue was dissociated by serial collagenase II (305 units mg^−1^ in HBSS, Gibco) digestions at 37 °C. CMs were purified from adherent cardiac cells via differential attachment, and any unattached cells after 1 h were considered enriched CMs. CMs were counted and seeded within 1–2 h following enrichment.

### Cell seeding and culture

Gel-Amin precursor solutions were prepared with 8% (w/v) GelMA, 3.5% (v/v) ChoA in complete culture medium (Dulbecco’s modified Eagle medium with l-glutamine supplemented with 10% (v/v) fetal bovine serum, 1% (v/v) penicillin-streptomycin) containing 0.5% (w/v) LAP. GelMA precursor solutions were prepared at 9.75% (w/v) in complete culture medium with 0.5% (w/v) LAP. Isolated CMs were pelleted and resuspended in either the Gel-Amin or GelMA precursor solution at a density of 7.5 × 10^4^ cells/μL. Encapsulated CM gel solutions were carefully loaded into the corresponding seeding port to form a liquid interface at the GelPin. This was repeated for both hydrogel groups, so they were both contacting the GelPin. The hydrogel solutions were then photo-crosslinked in situ with visible light (405 nm, 10 mW cm^2^) for 57 s (0.25 s μm^−1^ of hydrogel thickness). Following gelation, a complete culture medium was added to the chip. Cell cultures were incubated at 37 °C and 5% O_2_. Media exchanges occurred every two days.

### Live/Dead viability assay

A live/dead viability assay was conducted based on the manufacturer’s protocol (Thermo Fisher). To begin, glass slides were UV sterilized for five minutes on each side and plasma-treated for three minutes (Harrick Expanded Plasma Cleaner: PDC-002) to increase hydrophobicity. Hydrogels and CMs were prepared before seeding. CMs were encapsulated in either the Gel-Amin or GelMA hydrogel (n=6) at a density of 7.5 × 10^4^ cells/μL and seeded onto glass slides in a 12-well plate. Hydrogels were crosslinked, and the cells were cultured for up to seven days with media exchanges every three days. On days five and seven, post-seeding staining was performed. A separate 15 mL Falcon tube was filled with DPBS, 10 mL DPBS, 5 µL Calcein AM (0.5:1000), and 10 µL Ethidium Homodimer-1 (1:1000). Media was removed from the well plate, and 200 µL of staining solution was added to each well. The plates were incubated for 30 minutes at 37 °C. Following this, cells were imaged using a Zeiss inverted light microscope and analyzed using ImageJ.

### MPS fabrication

MPSs were fabricated and assembled as previously described.^9, 28^ Each layer was designed using 2D CAD, and a laser engraver (Epilog Zing 30W, Epilog Laser) was used to cut the designs into the materials as depicted in **Figure 1B**. The 3/16″ poly(methyl methacrylate) (PMMA, McMaster-Carr) upper layer featured a hollowed square to optimize media storage. The second layer was comprised of a 1/16″ PMMA sheet, which featured a central square to allow media to contact the membrane and cell cultures. Additionally, this layer featured two 1-mm-wide circular channels used as seeding ports for the polymer gel. This layer was sandwiched between 50-*μ*m-thick double-sided adhesive tape (966 adhesives, 3M, Maplewood, Minnesota). The third layer, a polycarbonate (PC) membrane with 1.0 μm diameter pores (Healthcare, Marlborough, MA), featured two circular through-holes with identical diameters to the previous layer. The membrane enabled fluidic access to the bottom channel, composed of cell cultures. The fourth layer was comprised of a 0.003″ polyethylene terephthalate (PET, McMaster-Carr) sandwiched between 50 μm double-sided adhesive tape. Before assembling the chips on the final layer, all chips were heat pressed for one minute at 56 °C. Following this, chips were then seeded onto the microelectrode array, where they were positioned to evenly separate the grid of electrodes. Post-assembly, the devices were stored under a vacuum at 37 °C for three to five days.

### MEA recordings

Electrophysiology was recorded using the multichannel system head stage (Multichannel Systems) based on a previously established protocol.^33^ To begin, the head stage was heated to 37°C using the TCX Controller application. Once this was achieved, the perimeter of the MEA (Multichannel Systems) was thoroughly wiped with a 100% ethanol solution to clean the electrodes. The MEA was then inserted into the head stage, and secondary containment was used to cover the MEA and prevent contamination. The MEAs were then left alone for ten minutes to bring CMs to a steady state. Recordings were collected using the software, MC_RACK. The recorded data was processed by applying a Butterworth 2nd-order filter. Spikes were determined using a threshold of -3 standard deviations of the peak-to-peak noise determined by the software. An electrode was not included in further analysis if the noise exceeded 1 mV. To evaluate signal propagating, a Synchronicity value, S, was found throughout the recording time (n=5). This value is obtained using the MATLAB-based SPIKY software^34^ by Thomas Kreuz and is standardized between a value of zero and one. A value of zero indicates a very synchronous culture where the spike is identical to others in the culture.

### Signal-to-noise ratio (SNR)

The SNR for the MEAs was calculated using the MC_RACK software (n=5). The background noise was measured for all electrodes at a randomized time interval. The determined spike amplitude was then recorded and inputted into Eq. 1 to evaluate the SNR, which evaluates voltage noise in units of decibels (dB):

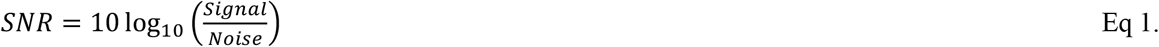

### Statistics

GraphPad Prism was used to assess statistical comparisons using a Mann-Whitney U test unless otherwise stated. For signal-to-noise ratio data, a lognormal distribution was assumed due to the logarithmic calculation (Eq. 1), and statistical analysis was performed using a lognormal Welch’s t test. Error bars on the graphs represent the standard error of the mean (SEM).*, **, ***, and **** represent a p value of <0.05, <0.01, <0.001, and <0.0001, respectively.

## Results

### Material Characterization

Both hydrogel formulations were previously mechanically and electrically characterized by Neuman *et al*.^27^ The mechanical properties of an 8% (w/v) GelMA hydrogel conjugated with 3.5% (v/v) ChoA(Gel-Amin) closely matched those of native cardiac tissue (10-15 kPa).^35^ A 9.75% (w/v) GelMA concentration was chosen as the control, as it has a comparable elastic modulus to Gel-Amin. Further, EIS was performed to evaluate the conductivity of both formulations, 123 × 10^−5^ S cm^−1^ and 375 × 10^−5^ S cm^-1^ were reported for GelMA and Gel-Amin, respectively. It is interesting to note that the conductivity of the Gel-Amin material is within the physiological range of the myocardium, while GelMA’s conductivity is below this limit.^19, 20^

### Ionically conductive hydrogels support high viability cultures for up to seven days

A viability assay was conducted to evaluate the capability of the 3D platform to support CM cell cultures. Gel-Amin was found to have a mean viability of 90.1% (± 6.94%) at five days post-seeding and that of 85.7% (± 4.95%) at seven days (**Fig. 2**). Additionally, the GelMA group had an overall viability of 75.3% (± 13.0%) and 78.4% (± 14.3%) on days five and seven, respectively. While there were no significant differences between the viability in the GelMA hydrogel compared to Gel-Amin, the Gel-Amin showed overall higher mean viability at five and seven days post-seeding. These results were consistent with Neuman et al.’s viability screen of Gel-Amin on peripheral neurons.^26, 27^

**Figure 2.**
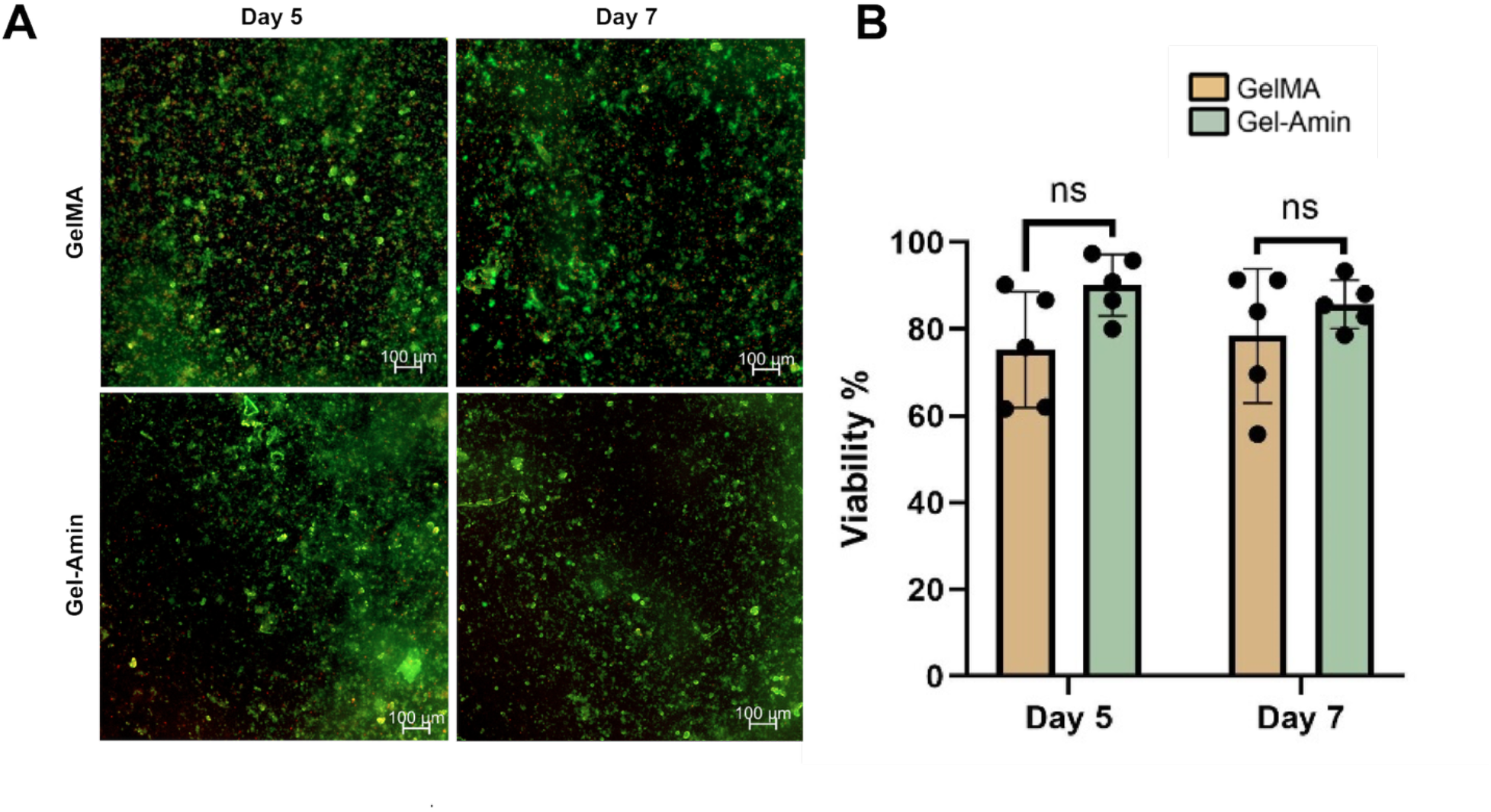
Live/Dead viability assay shows Gel-Amin and GelMA hydrogels support high viability cardiomyocyte cultures for seven days. A) Images captured on Zeiss Blue Inverted Light Microscope. Green dye indicates viable cells stained with Calcein AM. Red dye denotes dead cells. Scale bars = 100 µm. B) Cardiomyocyte viabilities were quantified using the analysis software, ImageJ. This software calculated the number of dead cells and live cells. Using the information, viability was found by dividing live cells by the total number of cells counted. No significant differences were observed between cardiomyocytes in the GelMA or Gel-Amin scaffold (n = 5, mean+/-S.D., p = 0.167).

### Chip fabrication and assembly

An MEA was used to evaluate the electrophysiology of the encapsulated CMs. A 3D MPS was designed to facilitate interfacing GelMA/Gel-Amin structures on the electrode footprint (**Fig. 1**). The MPS structure was bonded to commercially available MEAs with 3M 966 Transfer Tape. This design aims to place the encapsulated CMs positioned directly above the electrode grid of the MEA. Furthermore, the design was optimized with a GelPin to allow for dual seeding of the GelMA and Gel-Amin hydrogels on the same MEA platform. The chip underwent iterative designs to maximize media storage and an easy-to-seed compartment. No leaks were observed, and the MPS fit within the Multichannel Systems headstage (**Fig. 1**).

### Gel-Amin Amplifies Cardiomyocyte Signaling and Synchronicity

Using the previously described MEA Chip platform, the electrophysiology of CMs encapsulated in either the GelMA or Gel-Amin hydrogels was evaluated following seven days of culture. A raster plot of APs comparing GelMA and Gel-Amin exhibits clear spatial differences (**Fig. 3**). Qualitative analysis shows that the Gel-Amin environment promotes greater synchronicity in CM contractions. Further, a value of one denotes a unique spike. Throughout the time course of recording, the CMs in Gel-Amin a higher degree of synchronicity than GelMA CMs (**Fig. 3A**). The average synchronicity value for CMs in Gel-Amin (0.14) was significantly different than that of CMs in GelMA (0.27, p = 0.008, **Fig. 3B**). This result supports the theory that, due to the improved ionic conductivity, CM signals were amplified, which resulted in more interconnected tissue networks.

**Figure 3.**
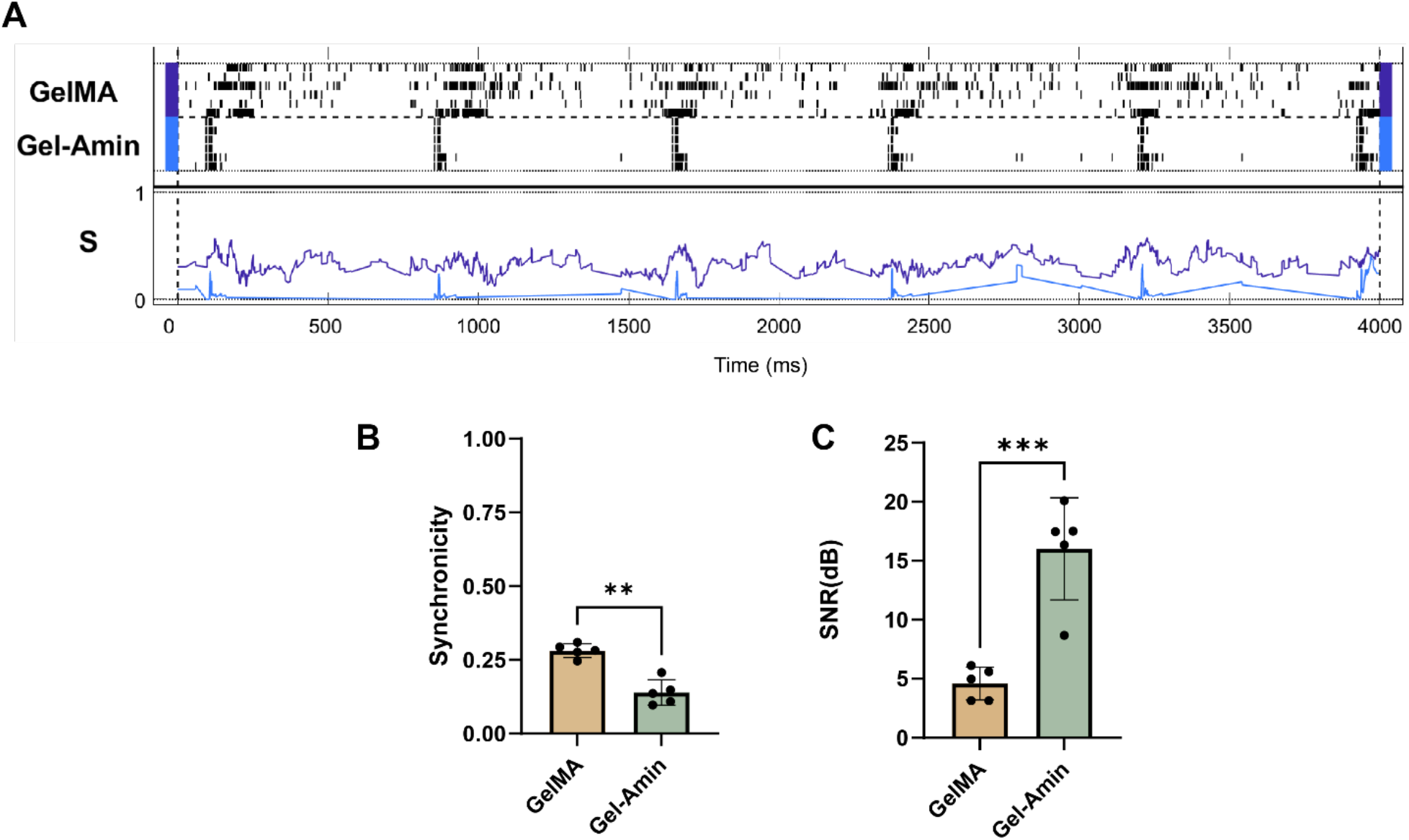
Evaluation of Cardiomyocyte Synchronicity. A) Raster plot shows spike timestamps from individual electrodes to evaluate the synchronicity (n=5). Spikes were determined from MEA electrophysiology data by applying a negative threshold of -3 standard deviations of the peak-to-peak noise determined by the software. Using the MATLAB application, SPIKY, the synchronicity (S) was determined by correlating the uniqueness of a spike between zero and one. Values near zero are considered identical to the other electrodes and would indicate strong synchronicity. Values near one denote a unique spike. B) Gel-Amin cultures are significantly more synchronous, improving the representation of native tissues. The average synchronicity value was calculated, and the values for each hydrogel group were compared. The results found that Gel-Amin had a significantly lower S value, indicating that the ionically conductive hydrogel improved biomimicry (p=0.008). C) Signal-to-noise ratio (SNR) comparison. The SNR ratio in the GelMA and Gel-Amin hydrogels was calculated by measuring the spike signal and noise and applying it to Equation 1. SNR across the electrodes was averaged for a mean SNR and compared to repeated experimental trials (n=5). The results found a 3-foldincrease in SNR in the Gel-Amin group compared to the GelMA control (p=0.0003). These results show that including a Bio-IL in the polymer network can improve conductivity and amplify cardiomyocyte signaling. This finding presents a novel strategy for 3D culture to record real-time, accurate electrophysiology data to evaluate the biological activity of *in vitro cell* cultures in a biologically relevant model.mean+/-S.D.

### The inclusion of ChoA significantly increases the signal-to-noise ratio (SNR)

The SNR was evaluated to determine the potential for signal amplification in the ionically conductive hydrogel. The average SNR for Gel-Amin was 16.0, and that of GelMA was 4.6 (**Fig. 3C**). It was found that the increased SNR was significant (p = 0.0003).

## Discussion

The cardiac conduction system’s architecture and electrical properties facilitate coordinated contraction from the nodes, through the atria, and descending the ventricles, yielding a high degree of pump efficiency.^36^ Changes in this structure and electrical function can lead to arrhythmias.^37^ To date, there are no *in vitro* models tackling changes in signal propagation via patterned biomaterials of different electrical properties.^38, 39^ Here, we present critical steps towards better modeling the cardiac system in a dish.

In this study, we aimed to evaluate an ionically conductive hydrogel for cardiac tissue engineering. We investigated its ability to enhance CM signaling, thereby developing a biomimetic cardiac model that can be adapted for drug development. There is an unmet need for biocompatible and conductive materials to generate *in vitro* models of excitable tissues to bridge the biological gap between preclinical and clinical settings. Our 8% (w/v) GelMA + 3.5% (v/v) ChoA (Gel-Amin) material showed enhanced ionic conductivity similar to native myocardium, optical transparency, and supported CM cultures with high viability. These results agreed with our previous work exploring GelMA + ChoA hydrogel formulations.^26,27^ Furthermore, we demonstrated that by crosslinking ChoA into the GelMA hydrogel, the signaling of encapsulated CMs is enhanced, allowing for electrophysiological recording via a simple planar MEA. These results suggest that hydrogels functionalized with Bio-IL present a novel strategy to overcome the innate non-conductive properties of many hydrogel polymers while retaining biocompatibility.

Although our Gel-Amin hydrogel shows promise, extensive work has been done to overcome the inherent non-conductivity of biomaterials for biomedical applications. One strategy involves the addition of nanoparticles like graphene or carbon nanotubes to develop conductive composites.^41, 42^ These materials have significantly improved electrical conductivity and have high mechanical strength.^43^ Shin et al., showed that a reduced graphene oxide (rGO) and GelMA composite biomaterial showed increased electrical conductivity and mechanical strength, which improved the spontaneous beating of primary CMs.^44^ Another strategy to improve the electrical properties of a biomaterial involves synthesizing polymers with increased conductivity, like polypyrrole (PPy) or poly(3,4-ethylenedioxythiophene) (PEDOT).^45^ Extensive research into conductive polymers allows for favorable electrical properties in a biomaterial scaffold. However, both strategies face significant limitations for cardiac tissue engineering, including cytotoxicity, genotoxicity, biodegradability complications, difficult synthesis methods, and limited optical transparency.

Functional assessment of electrophysiology in 3D cultures has presented technical challenges due to hydrogel non-conductivity. Creating interfacing 3D structures of varying geometries is a key advantage of our layer-by-layer MPS fabrication techniques.^9,40^ Real-time functional measurements offer clear advantages over terminal characterization such as label-free monitoring and provides information that may not be observable. We hypothesized that the inclusion of ChoA would amplify cardiomyocyte signaling due to increases in the ionic conductivity of the biomaterial. To investigate this, a MPS was designed to evaluate the ability to monitor CM activity simultaneously. Our hypothesis was supported as we discovered that incorporating ChoA into the GelMA polymer network resulted in amplified CM signaling. The SNR for the Gel-Amin group was significantly greater than the control group (**Fig. 3**). Analysis of beating patterns via a raster plot and synchronicity assessment demonstrated that CMs encapsulated in Gel-Amin had a more united network, which led to synchronous beating. Coordinated contraction is pivotal to cardiac pump function. In the future, this design could be adapted to design architectures to geometrically define conduction waves and drug screening to monitor changes in cardiac output in a real-time, label-free, and cost-effective manner.

While the Gel-Amin hydrogel and MPS demonstrated promising results in amplifying CM signaling, several limitations must be acknowledged. First, although ChoA incorporation improved ionic conductivity, the long-term stability and degradation profile of the hydrogel under culture conditions remain unknown. Another limitation is the use of neonatal rat primary CMs for functional assessments. However, the overarching goal of this system is to emulate the human cardiac system. Future work could adapt these methodologies using humanized cardiomyocyte models, such as human induced pluripotent stem cell-derived cardiomyocytes (hiPSC-CMs). Incorporating these cells would enhance the biomimicry of the device and support the development of a mature, humanized MPS to study tissue-specific behavior *in vitro*. Additionally, this study does not present data validating the platform for drug screening applications. To address this, future experiments could adapt the MPS to evaluate various cardiac-modulating compounds (e.g., epinephrine, isoproterenol) and record the resulting electrophysiological changes. Such validation would be critical to establishing the platform’s utility in preclinical drug development and cardiotoxicity screening.

There is a need for improved *in vitro* models to bridge the gap between preclinical and clinical settings to improve FDA approval rates by predicting drug activity. The development of this MPS presented a high-throughput, cost-effective screening device to monitor CM signaling networks. These results provide a framework for *in vitro* cardiac models that could be applied to predict potential cardiotoxicity for patients to improve the success rates of drugs. The adoption of systems like this by pharmaceutical companies would help reduce drug project costs and time spans and ultimately increase the availability and affordability of life-saving therapeutics for patients.

## Author Contributions

D.P., K.N., A.K., and R.K. conceived the project. D.P. performed all the cell and MEA experiments, and K.N. performed the material characterization experiments. D.P. and N.B. analyzed data under the advice of R.K. D.P., A.K., and R.K. prepared the figures and wrote the article. K.N. and N.B. edited the article.

## Acknowledgements

The authors thank the Department of Chemical Engineering for support and Rob Egan for technical machining assistance. We would also like to thank Kyla Nichols for her assistance in the training with the microelectrode system and William Doherty for his help with the isolation and dissociation of primary cells. We thank Dr. Joshua Gallaway and Benjamin Leiffer for their assistance with the VSP Potentiostat. The authors would like to acknowledge funding for this project through Northeastern PEAK research fellowships, NIH NIGMS, and NASA.

